# Reduction of chromosomal instability and inflammation is a common aspect of adaptation to aneuploidy

**DOI:** 10.1101/2023.12.22.572964

**Authors:** Dorine C. Hintzen, Michael Schubert, Mar Soto, René H. Medema, Jonne A. Raaijmakers

## Abstract

Aneuploidy, while detrimental to untransformed cells, is notably prevalent in cancer cells. This indicates that cancer cells have the ability to surmount the initial stress responses associated with aneuploidy, enabling rapid proliferation despite aberrant karyotypes. To generate more insight into key processes and requirements underlying the adaptation to aneuploidy, we generated a panel of aneuploid clones in p53-deficient RPE-1 cells and studied their behavior over time. As expected, *de novo* generated aneuploid clones initially displayed reduced fitness, enhanced levels of chromosomal instability and an upregulated inflammatory response. Intriguingly, after a prolonged period of culturing, aneuploid clones exhibited increased proliferation rates while maintaining aberrant karyotypes, indicative of an adaptive response to the aneuploid state. Interestingly, all adapted clones displayed reduced chromosomal instability (CIN) and reduced inflammatory signaling, suggesting that these are common aspects of adaptation to aneuploidy. Collectively, our data suggests that CIN and concomitant inflammation are key processes that require correction to allow for fast growth. Finally, we provide evidence that amplification of oncogenic KRAS can promote adaptation.

## Introduction

Each time a cell divides it must distribute its genomic content equally amongst daughter cells. Cell division is a highly regulated process, safeguarded by various checkpoints such as the spindle assembly checkpoint (1). However, when errors occur, chromosomes can missegregate, giving rise to cells with erroneous karyotypes. The state of having an aberrant number of chromosomes is known as aneuploidy.

Aneuploidy manifests deleterious effects at both the organismal and cellular levels, resulting in developmental defects and reduced cellular proliferation (2–8). The detrimental effects of aneuploidy have been attributed to imbalances in coding genetic material (4,9). It is widely acknowledged that aneuploidy results in the altered expression of the genes located on the involved chromosome(s). While RNA expression largely scales with DNA copy number (4–6,10–14), large-scale dosage compensation is observed at the protein level in an attempt to maintain complex stoichiometry and appropriate levels of dosage sensitive factors (6,10,13,15–17). The disruption of proteostasis upon aneuploidy results in an increased burden on protein synthesis, folding and degradation machineries, often leading to proteotoxic stress (18). Indeed, in both aneuploid yeast and human cell models, indications of proteotoxic stress and a higher dependency on processes related to the maintenance of proteostasis have been extensively documented (6,7,13,19–24).

Aneuploidy has also been linked to enhanced genetic instability (4,25) and replication stress (4,23,26–29), which can be caused by imbalances in crucial protein complexes (26). Besides, numerous studies have reported metabolic alterations in response to aneuploidy (4,6,30–33). These metabolic alterations might be caused by the enhanced energy requirements to deal with the challenges induced by the aneuploid state. Importantly, the maintenance of metabolic homeostasis is also contingent on the stoichiometry of metabolic regulators and enzymes, making it susceptible to the effects of gene imbalances. Moreover, elevated levels of reactive oxygen species (ROS) are present in aneuploid cells (7,13,34,35), which poses a further threat to genomic stability. Altogether, aneuploidy has a severe impact on the proteostasis of the cell, thereby disrupting many cellular processes, inducing stress responses and forming a major threat to the fitness and genomic stability of the cells.

While aneuploidy arises as a consequence of errors during mitosis, aneuploidy has also been shown to be a driver of ongoing mitotic errors, a phenomenon termed chromosomal instability (CIN) (25,26,28,36–39). The induction of CIN has also been linked to gene imbalances (25,39), and recent work shows that CIN levels specifically correlate to the amount of gained coding genes (40). Continuous CIN can evoke an inflammatory response via the cGAS-STING pathway as a consequence of genetic material that ends up in the cytoplasm due to rupture of micronuclei or chromatin bridges that persist (41–44). Additionally, CIN can give rise to cellular populations harboring complex karyotypes, capable of inducing inflammation through entering a senescent-like state and the acquisition of a senescence-associated secretory phenotype (SASP) (29,45).

In light of the extent of stress responses elicited by aneuploidy, it is apparent that aneuploidy represents a highly detrimental condition. At first sight it therefore seems paradoxical that aneuploidy is highly prevalent in cancer, a disease characterized by fast and uncontrolled proliferation, with approximately 90% of all solid tumors being aneuploid (46–49). This phenomenon is therefore referred to as the “aneuploidy paradox” (50,51). Many studies have described the positive effects of aneuploidy and CIN on tumor heterogeneity, tumor progression and its contribution to therapy resistance (52–54), highlighting the benefits for tumors to induce and maintain aneuploidy. However, the adaptive mechanisms of tumor cells to the tolerance of such aberrant karyotypes are far less understood. Understanding this is important, as certain adaptation mechanisms could be exploited for therapy development.

One important factor in aneuploidy tolerance in human cells is p53 (55,56). However, even in the context of p53-deficiency, *de novo* aneuploidies severely impact cellular fitness (25,26,40,46,57). A few tolerance mechanisms have been shown to be at play independently of p53. For example, in yeast, that do not carry a TP53 gene, adaptation to aneuploidy has been shown to involve mutations in genes involved in mTOR signaling (58) or in protein degradation (12). Initial tolerance to *de novo* aneuploidies in human cells has been shown to be determined by p38-mediated metabolic alterations (59). Besides, loss of BRG1, a member of the SWI/SNF complex, has been shown to alter initial aneuploidy tolerance, possibly also by affecting the metabolic state of the cell (60). However, less is understood about the processes underlying long-term adaptation to aneuploidy in human cells. One study shows that restoring Hsp90-mediated protein folding promotes adaptation to aneuploidy in cell culture (19). However, the exact adaptive mechanisms that are employed by aneuploid cells are currently not clear.

In this study, we set out to elucidate the general requirements and mechanisms via which human cells adapt to aneuploidy over a prolonged period of time. Because of the central role of p53 in aneuploidy tolerance (55), we made use of a near-diploid p53-deficient human cell line. Also, to exclude karyotype-specific adaptation pathways, we make use of a large panel of different aneuploid clones with a variety of aneuploidy levels. Using this system, we systematically uncovered commonalities intrinsic to cells adapting to aneuploidy.

## Results

### Cells can adapt to aneuploidy

To investigate mechanisms of adaptation to aneuploidy, we generated a panel of aneuploid clones originating from an hTERT-immortalized retinal pigment epithelial cell line (RPE-1). Notably, given the pivotal role of p53 in aneuploidy tolerance, we conducted these experiments using clones harboring a stable shRNA-mediated knockdown of p53 (55). The generation of *de novo* aneuploid clones was described previously (40). We selected 14 clones with a variety of aneuploidy levels, mostly involving entire chromosomes (Fig S1A). We cultured these clones for a time span of several months. To account for potential effects due to extended culturing as well as aging, we subjected the parental cells to the same prolonged culturing conditions. At monthly intervals, we measured the doubling times of these clones using live-cell imaging as a read-out for cellular fitness and, by extension, adaptation (see Fig.1A for the working pipeline). As expected, and in line with our previous report (40), all clones initially displayed a growth defect, albeit to different extents. Remarkably, all selected clones exhibited an increase in proliferative capacity over time, indicative of adaptation (Fig. 1B).

**Figure 1:**
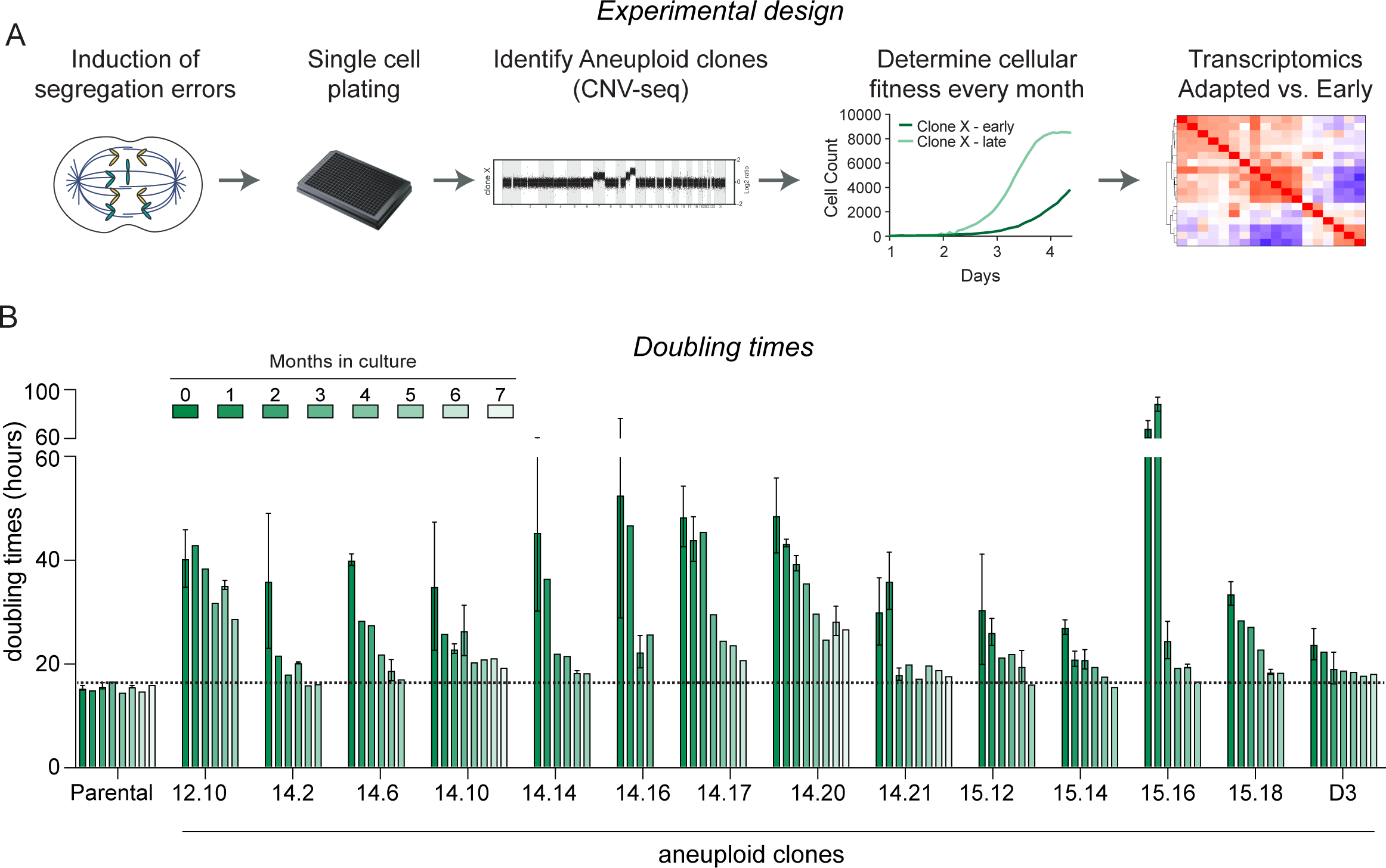
Cells adapt to aneuploidy A. Working pipeline. Parental RPE-1 p53KD cells were treated with MPS1 and CENP-E inhibitors to induce segregation errors and plated single cell to generate clones. The ploidy of the clones was determined using CNV sequencing and only aneuploid clones were kept for further investigation. Cells were subjected to long-term culturing and adaptation was monitored using live-cell imaging. To elucidate adaptation mechanisms, RNA sequencing was performed. B. CNV plots of clone 14.16 comparing copy number alterations early and adapted. C. Doubling times of parental RPE-1 p53KD, and aneuploid clones determined via live-cell imaging performed at monthly intervals. Error bars indicate standard deviation. In-between time points were only imaged once and hence do not contain error-bars.

### Full karyotype correction is not a common mechanism of adaptation

To gain insights into the adaptation process of the adapted clones, we conducted RNA sequencing on all 14 early aneuploid clones and their adapted counterparts. For the adapted clones, we selected the first time point at which they proliferated at their maximum capacity, minimizing the potential confounding effects of aging. The most obvious path to adaptation would be karyotype correction, where aneuploid clones revert to a (more) euploid karyotype. Since RNA levels largely scale with DNA copy number (4–6,10–14), we first utilized the transcriptome data to extract karyotype information (Fig. 2A: red regions reflect gains, blue regions reflect losses). This method confirmed the karyotypes of the early clones determined with CNV-seq data, validating the reliability of using RNAseq reads for deducing karyotypes (compare Fig. S1A and Fig. 2A). For one clone we determined that upon adaptation the RNAseq reads still reflect those of DNA copy number (Fig. S1B). This not only shows that the deduction of karyotypes can be reliably performed both in early and adapted clones but it also suggests that large-scale dosage compensation on the level of the transcriptome is unlikely to contribute to adaptation. When we examined the karyotypes of early clones in comparison to their adapted counterparts, we observed distinct patterns of karyotype alterations. Importantly, although some clones displayed a relatively stable karyotype upon adaptation (ie. 14.16, 14.20), most clones displayed some levels of karyotype alteration. To obtain a quantitative measurement of these alterations, we determined the number of imbalanced genes for each clone by applying cutoffs on the fold change values of gene expression (see methods). Using these measurements, we categorized the karyotype progression upon adaptation into four distinct groups: “more complex” (karyotypes obtain more imbalanced genes), “evolved” (karyotypes changed but maintained a highly aneuploid karyotype), “simplified” (remains aneuploid, but less complex karyotypes), and “reverted” (karyotype similar to the parental RPE-1).

**Figure 2:**
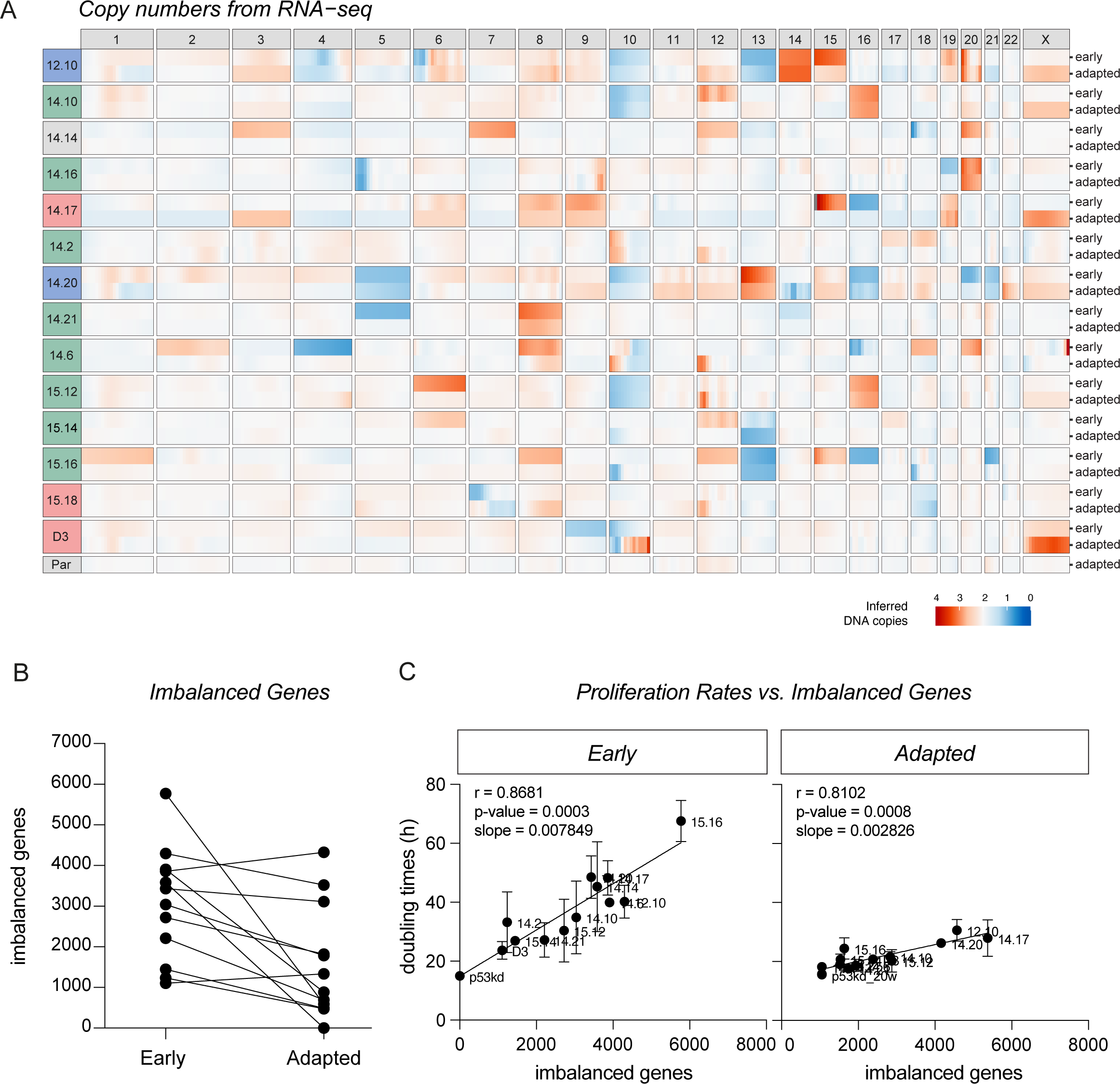
Full karyotype correction is not a common mechanism of adaptation A. Karyotypes of early and adapted clones derived from RNA sequencing data, using RPE-1 parental p53KD cells as a reference. Clones are color coded by their adaptation behavior: blue clones are clones with an “evolved” karyotype, green clones belong to the “simplified” karyotypes class, grey clones refer to clones with a completely “reverted” karyotype and red clones obtained “more complex” karyotypes. B. Number of imbalanced genes per clones calculated from RNA sequencing data in early and adapted clones. Lines connect early clones to their adapted counterparts. C. Spearman correlation between total coding-gene imbalances per clone and proliferation rates measured in Fig. 1B for early and adapted clones. Error bars indicate standard deviation. Line shows linear regression model fitted to the data.

Out of 14 clones analyzed, a subset of clones displayed an “evolved” karyotype (14.3%) or became even “more complex” (21.4%) in terms of their karyotypes. However, the majority of clones exhibited karyotype simplification (57.1%). Notably, only one clone (14.14) fully reverted its karyotype to the parental karyotype, emphasizing that complete karyotype correction is not a common mechanism of adaptation. Importantly, when we compared the number of imbalanced genes of all early clones to their adapted counterparts, we observed that the extent of karyotype simplification is relatively modest for most clones (Fig. 2B). This suggests that the observed karyotype simplification cannot fully account for the enhanced proliferation rates, that often reaches the rates of parental RPE-1 cells.

It has been shown by us and others that proliferation rates strongly correlate to the number of imbalanced genes (4,6,10,13,40). If adaptation to aneuploidy is indeed not solely driven by karyotype simplification but other processes contribute to the adaptation process, we predicted that the impact of imbalanced genes on proliferation rates will be less pronounced in adapted clones as compared to their early counterparts. Consistent with our earlier investigation, we observed that imbalanced genes exhibit a strong correlation with doubling times in early clones (Fig. 2C, r=0.8681, p=0.0003), suggesting that the extent of gene imbalances indeed determines the extent of the proliferation defect, at least in the early clones. In line with our previous observations, the number of imbalanced genes explained the proliferation defect better as compared to the number of gained or lost genes separately (Fig S1C). Importantly, upon adaptation, we observed a strong reduction in the slope of the linear regression model applied to imbalanced genes’ data in the adapted clones (Fig. 2B, shifting from 0.007849 to 0.002826), in agreement with our prediction. This suggests that the impact of gene imbalances on doubling time reduces during the adaptation process, resulting in enhanced proliferation rates. It is important to note that the slope does not reach 0, underscoring that even in adapted clones, the presence of imbalanced genes continues to exert an effect on proliferation, albeit to a lesser degree compared to early clones. Altogether this data shows that karyotype simplification or correction is not a common driver of adaptation but that active adaptation mechanisms must be at play.

### Reduced CIN and inflammation are common aspects of adaptation

Adaptation may involve the development of mechanisms to downregulate the initial stress responses triggered by aneuploidy or to improve the ability to tolerate these stress responses. To gain insight into the underlying mechanisms through which clones adapt to aneuploidy, we performed RNA sequencing and executed gene set enrichment analysis (GSEA) to assess which hallmarks and which biological processes were mostly altered during adaptation (Fig. 3A, Fig. S2A). Hallmarks that were most prominently upregulated upon adaptation were hallmarks associated with cell cycle, such as E2F targets and G2-M checkpoint (Fig. 3A, B), which is expected, as the adapted clones have increased proliferation rates (Fig. 1B). Similarly, predominantly upregulated biological processes are also attributable to the increased cellular growth such as translation, ribosome biogenesis, mRNA processing, and DNA replication (Fig S2A). Hallmarks that were most prominently downregulated upon adaptation were hallmarks associated with inflammatory responses such as TNFα signaling via NF-κB and IL-6/JAK/STAT3 (Fig. 3A and B). Importantly, these alterations were reproducibly observed in the majority of individual clones (Fig. 3B and Fig. S2B, C), suggesting that these are uniform responses in adaptation. Importantly, most alterations were in fact corrections of the early response to aneuploidy (Fig. 3A and Fig. S2A, see middle row). When we specifically filtered on alterations unique to the adapted clones, which were unaltered in early clones, we only found peroxisomes and glycolysis to be specifically upregulated (Fig. 3C). This suggests that adapted clones undergo metabolic rewiring.

**Figure 3:**
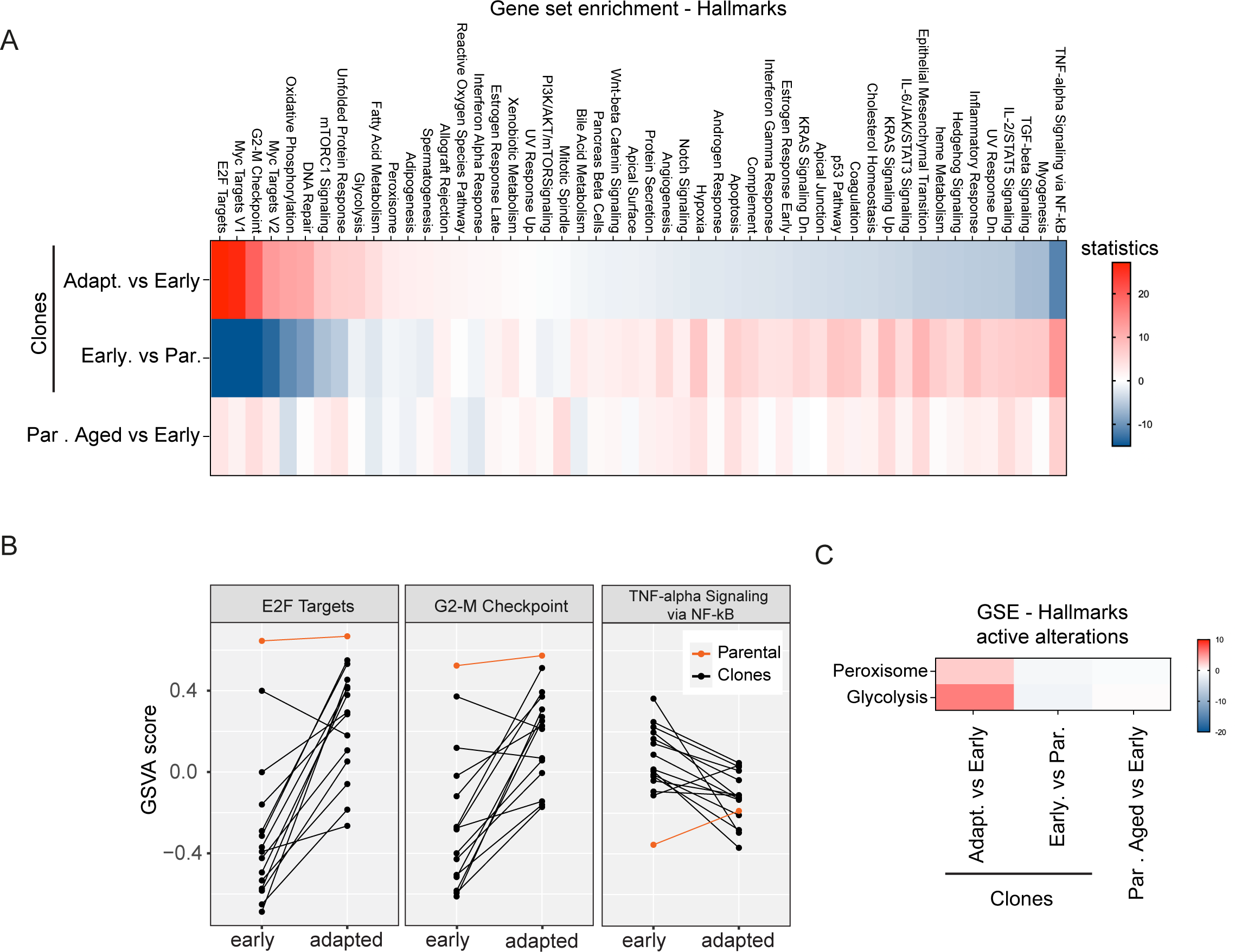
Reduction of inflammation and upregulation of cell cycle hallmarks are common aspects of adaptation A. Gene set enrichment for hallmarks based on the transcriptional alterations observed in adapted clones compared to their early counterparts, colors indicate Wald statistics. The top 50 most altered gene sets in the adapted clones are displayed (top row). Alterations in early clones over parental (middle row) and aged parentals (bottom row) are displayed as a reference. B. Plots showing Gene Set Variation Analysis (GSVA) for a selected number of hallmarks that show high alteration upon adaptation for the individual clones. Lines connect early clones with their adapted counterparts. The parental cells are displayed in orange as a reference. C. Selected hallmarks (panel A) that showed alterations upon adaptation but that were minimally altered in early clones versus parental cells, indicative of pathway activation rather than correction.

It is known that aneuploidy itself can drive CIN and CIN is an established instigator of inflammation. Therefore, we hypothesized that the reduction in inflammation might be a consequence of decreased aneuploidy-induced CIN during the process of adaptation. To test this, we conducted live-cell imaging on both early aneuploid clones and adapted clones and quantified the frequency of errors that occurred during anaphase. As expected, we observed that all early clones showed elevated CIN levels compared to parental cells (Fig. 4A). Interestingly, adapted clones all displayed lower levels of CIN when compared to their early counterparts, suggesting that downregulation of CIN is a common aspect of adaptation. Indeed, we find that the reduced CIN rates strongly correlate to the improved proliferation rates upon full adaptation (Fig. 4B). To test if CIN levels follow the improved proliferation rates also during the adaptation process, we monitored missegregation rates over the time course of the adaptation process for several clones (Fig. S3A). We observed that CIN levels consistently decreased during the adaptation trajectory in all clones, albeit with different kinetics. Overall, the decrease in CIN resembled the adaptation behavior in terms of doubling times which we exemplified by plotting doubling times against CIN rates, where both CIN levels and doubling times decrease simultaneously during adaptation (Fig. 4C, D).

**Figure 4:**
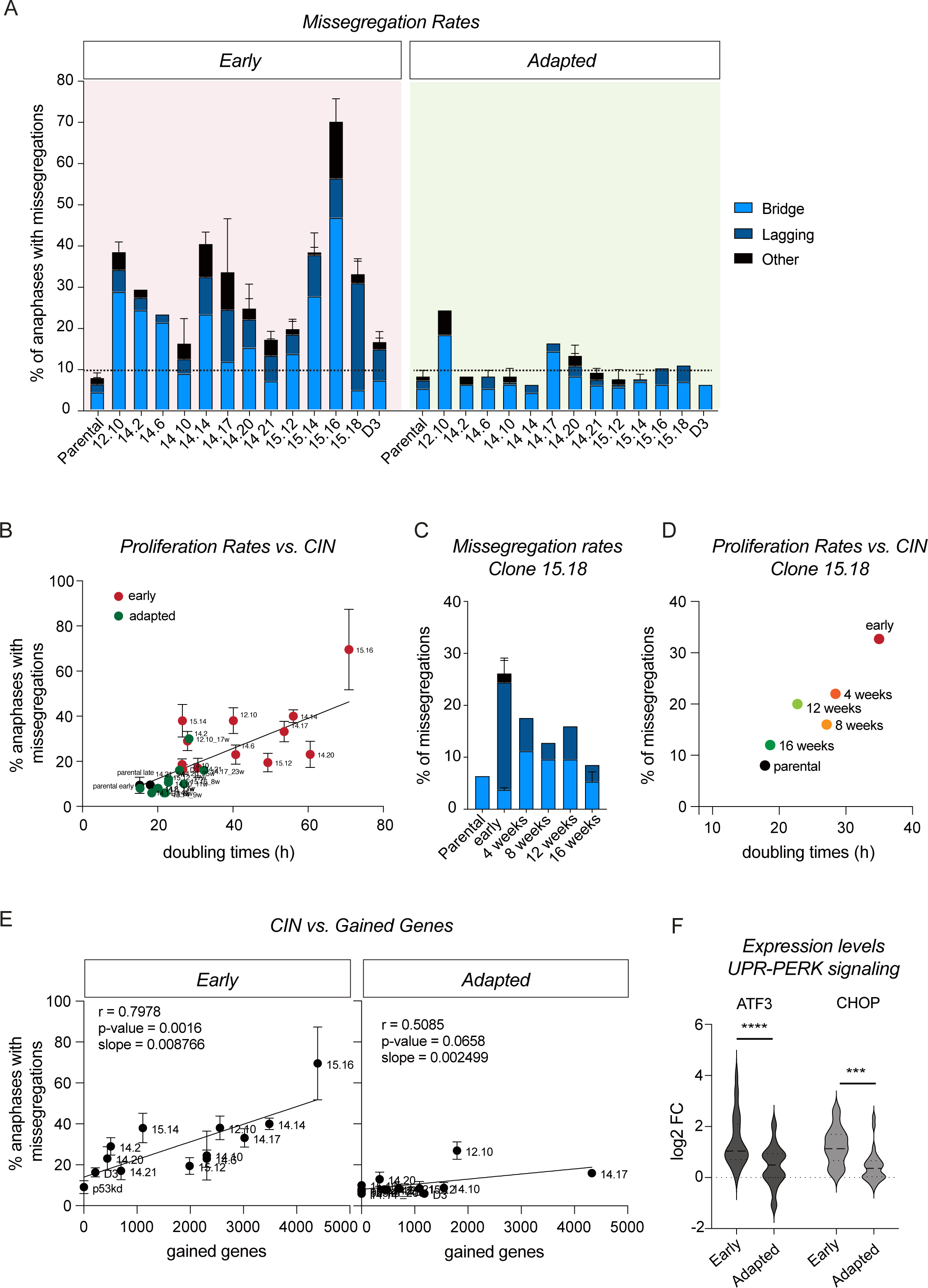
Reduction of CIN is a common aspect of cells adapting to aneuploidy A. Chromosome missegregation rates determined by live cell imaging of parental RPE-1 p53KD cells and aneuploid clones from Fig. S1A, divided into three subcategories: lagging chromosomes, anaphase bridges and others (multipolar spindle, polar chromosome, cytokinesis failure, binucleated cells). Bars are averages of at least 2 experiments and a minimum of 50 anaphases were analyzed per clone per experiment. Error bars indicate standard deviation. B. Correlation between the proliferation rates as measured in Fig. 1A and the missegregation rates as a percentage of anaphases as measured in A. C. Chromosome missegregation rates measured with intervals of a month for clone 15.18. D. Correlation between proliferation rates as measured in Figure 1A and missegregation rates as a percentage of anaphases as measured in C for clone 15.18. E. Spearman correlation between the number of RNA-sequencing derived gained genes per clone and the level of CIN as determined in A. Error bars indicate standard deviation. F. Violin plots showing the expression levels of ATF3 and CHOP in early and adapted clones extracted from the transcriptome data. An ordinary two-way ANOVA was performed between parental and clones. P-values are assigned according to GraphPad standard.

In our previous research, we demonstrated that the gained genetic material is the main determinant of CIN (40). In accordance, we observed the strongest correlation between gained genes and missegregation rates in early clones in this study (r = 0.7978, p = 0.0016) (Fig. 4E and Fig S3B). Interestingly, upon adaptation we observed a profound decrease in the slope of the correlation between gained genes and CIN levels (0.008766 to 0.002499), suggesting that gained genetic material has a dramatically lower impact on CIN levels in adapted clones as compared to early clones.

It was shown before that proteotoxic stress can be a driver of CIN (40). We found that the early aneuploid clones express higher levels of the two key players downstream of PERK activation, namely ATF3 and CHOP/GADD153 (61) (Fig. 4F), suggesting that these clones indeed experience proteotoxic stress, which could explain their increased CIN levels. Furthermore, upon adaptation, both ATF3 and CHOP were significantly downregulated, nearly reaching the levels of parental cells (Fig. 4F), suggesting that proteotoxic stress is no longer induced, despite the presence of abnormal karyotypes. These data suggest that adapted clones reduced the PERK-associated unfolded protein response, which could be responsible for the reduced CIN levels.

### Multiple consequences of CIN contribute to inflammation

Our study shows that aneuploidy triggers CIN and inflammation in early clones. Since both CIN and inflammation are corrected simultaneously upon adaptation, we speculate that these are directly linked and their correction might be required for adaptation. To uncover potential drivers of inflammation in the early clones, we performed GSEA on the transcriptome data. Gene sets for STING signaling, NF-κB activation and SASP were all positively enriched in the early clones (Fig S4A), indicating that multiple inflammatory pathways are activated. To investigate if the inflammatory response is a direct consequence of CIN, we aimed to compare the inflammatory response in our early clones to that of parental cells with induced missegregations. To this end, we treated RPE-1 p53KD parental cells with CENP-E and MPS1 inhibitors for 48 hours to induce missegregations and performed transcriptomics. Strikingly, we found very similar enrichment scores for the different inflammatory gene sets upon acute CIN compared to early clones. This suggests that the inflammatory response in early aneuploid clones can be largely explained by the enhanced CIN levels. Moreover, it shows that this inflammatory response is complex and involves multiple pathways. To explore the impact of an upregulated inflammatory response on cell proliferation, we utilized synthetic cGAMP to mimic the activation of the cGAS-STING pathway in non-aneuploid parental cells. Although earlier reports described the absence of cGAS and STING from RPE-1 cells (62), we found that STING is expressed in our RPE-1 cells (Fig 5C). Most importantly, we found that the cGAS/STING pathway can be activated in RPE-1 cells as we observed downstream target cytokine expression using different concentrations of cGAMP (Fig. 5A). We selected the highest dose for further experiments, as all tested cytokines were upregulated, indicating the successful activation of an inflammatory response. Importantly, this activation was at least in part dependent on STING as siRNA-mediated depletion of STING suppressed the induction of cytokines induced by cGAMP (Fig. S4B). Next, we assessed cellular fitness by measuring proliferation rates. Interestingly, we observed that parental cells treated with cGAMP have a higher doubling time (Fig. 5B and Fig. S4C), indicating that inflammation can result in decreased proliferation. To evaluate if reducing cGAS/STING-mediated inflammation in early clones could rescue the slow growth phenotype, we aimed to suppress cGAS-STING signaling via STING depletion. However, successful downregulation of STING did not result in increased proliferation rates in early clones. These findings indicate that the CIN-induced inflammation observed in early clones is multifactorial and although there is evidence for STING-signaling (Fig S4A), activation of cGAS/STING does not have a dominant contribution to the slow growth phenotype.

**Figure 5:**
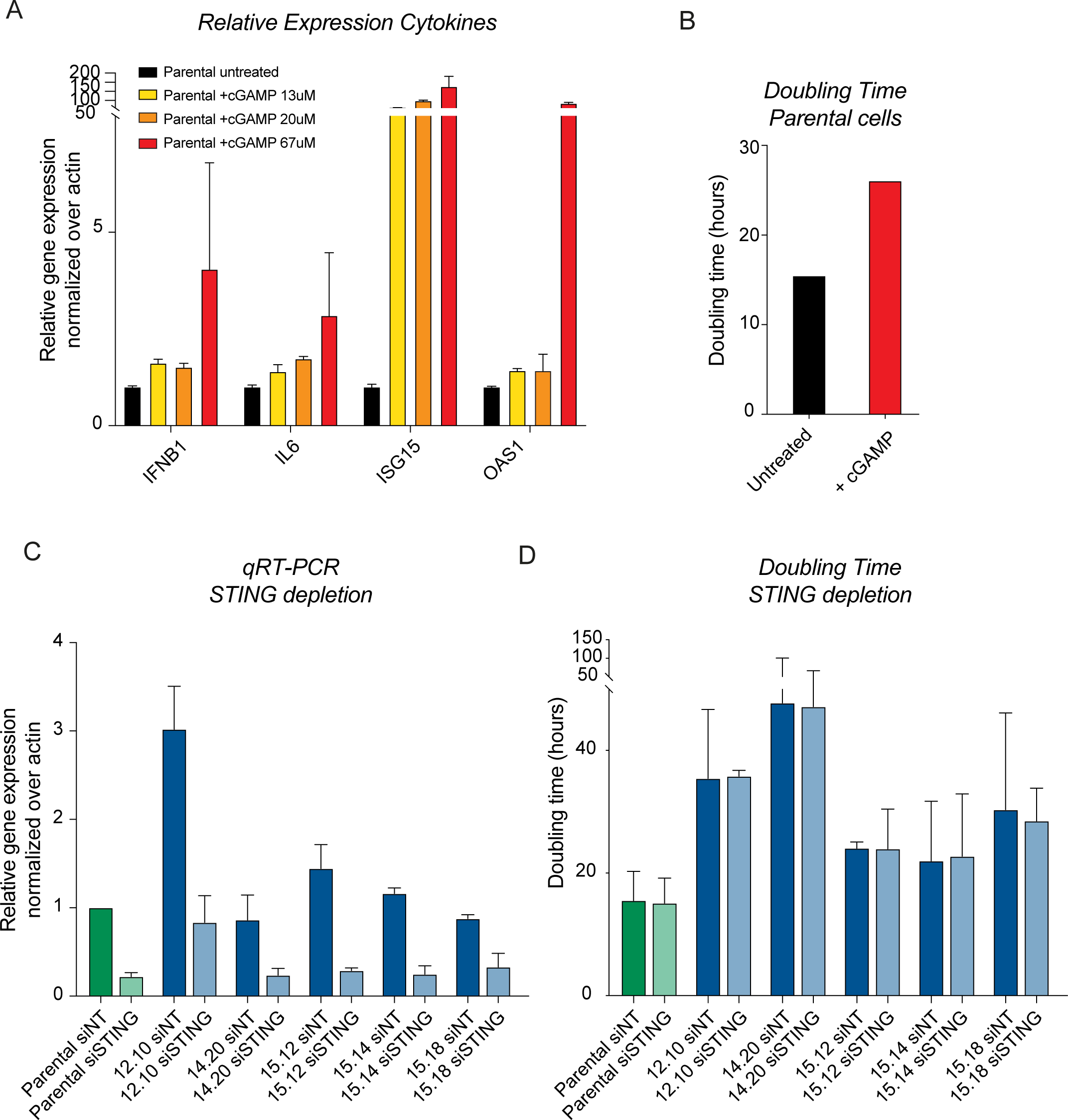
Investigating the impact of cGAS-STING signaling on proliferation A. mRNA levels of inflammatory response cytokines determined by qRT-PCR of parental RPE-1 p53KD, untreated or treated for 24 hours with different doses of cGAMP. Values were normalized to actin and are displayed relative to expression levels in untreated parental cells. Bars show mean expression levels; error bars indicate upper and lower limits. B. Average doubling times of parental RPE-1 p53KD untreated and treated with 67μM cGAMP added 2 hours prior to imaging, determined by live-cell imaging. C. mRNA levels of STING determined by qRT-PCR of parental RPE-1 p53KD and early aneuploid clones treated with siNT or siSTING. Values were normalized to actin and are displayed relative to expression levels in siNT parental cells. Bars show mean expression levels; error bars indicate standard deviation between two experiments. D. Doubling times of parental RPE-1 p53KD and early aneuploid clones treated with siNT or siSTING, determined via live-cell imaging. Two independent experiments were performed. Error bars indicate standard deviation.

### Amplification of mutant KRAS can drive adaptation

We established that reduced CIN and reduced inflammation are common aspects of adaptation to aneuploidy. We next set out to identify potential drivers of this adaptive behavior. Although we excluded karyotype simplification as a driver, karyotype alterations were commonly observed upon adaptation (Fig. 2 A, B). Previous studies suggested that specific aneuploidy patterns might arise to provide survival benefits (46,63–66). In order to determine whether our clones exhibit specific aneuploidy patterns during adaptation, we examined the karyotype alterations in more detail. Strikingly, we found that 4 out of the 14 clones displayed a chromosome 12p amplification upon adaptation (Fig 2A, clone 14.2, 14.6, 15.12 and 15.18). Moreover, 2 clones displayed an entire chromosome 12 amplification (clone 12.10 ad 14.20).

Importantly, one well-known oncogene, KRAS is located on 12p and one of the alleles of RPE-1 cells is mutated to an oncogenic version of KRAS (67). Moreover, the mutant allele of KRAS is amplified from 1 to 2 copies in parental RPE-1 cells (See micro-amplification in Fig S1A on chr12p and (67). We explored the possibility that acquiring an extra copy of the oncogenic KRAS allele provides a survival benefit to aneuploid clones. If this is true, we predict that the gained alleles consistently involve the mutant allele and not the WT. For this we determined the relative DNA copy number of the WT and mutant alleles in parental cells as well as two of the clones displaying 12p amplification (15.12 and 15.18) and 1 clone that displayed an entire 12 amplification (12.10), using TIDE (68). As parental cells carry 1 WT allele and 2 mutant alleles, the expected mutant/WT ratio is 0.66. Indeed, while parental cells and early clones display the expected ratio of mutant/WT alleles, the adapted clones all display enhanced copy number of the mutant allele, suggesting that indeed the allele containing oncogenic KRAS is amplified in the adapted clones with a 12p or 12 amplifications (Fig 6A). This increased copy number also leads to enhanced expression of the mutant KRAS (Fig. 6B).

**Figure 6:**
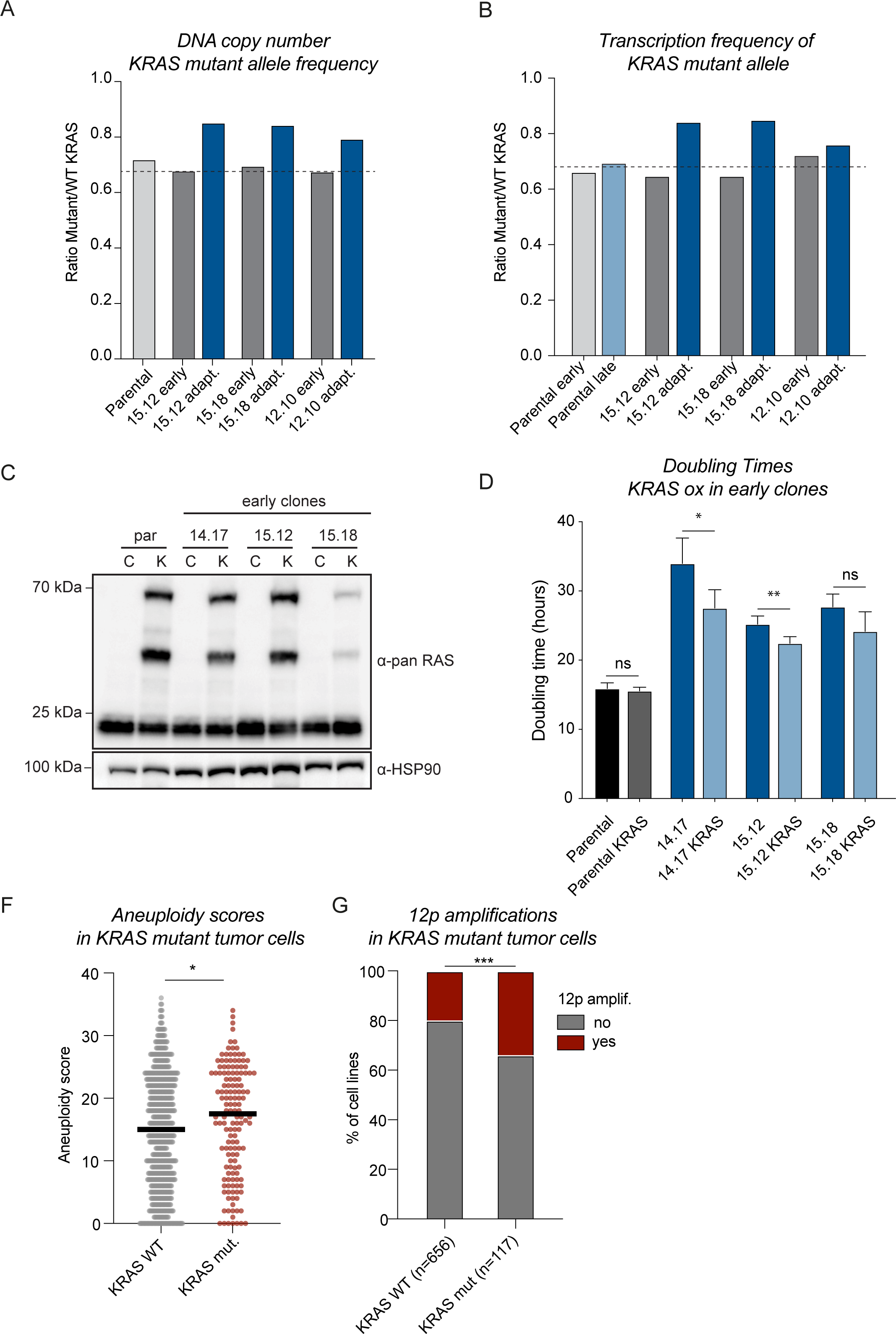
Overexpression of mutant KRAS can drive adaptation A. TIDE analysis was done on isolated DNA from parental RPE-1 cells and three clones (early and adapted). As a reference DNA isolated from Hela cells was used as HeLa cells have no described KRAS mutations. The graph depicts the relative frequency of the mutant allele over the WT allele. A consistent increase in mutant allele frequency is observed upon adaptation in the 3 selected clones. B. Reads from the transcriptome analysis were categorized as ‘mutant’ or ‘WT’ reads based on the presence of the absence of the known KRAS mutation. Only reads were quantified that span the mutant sequence and reads that could not be classified with certainty were excluded from the analysis. C. Western blot analysis shows successful overexpression of KRAS in the selected early clones and parental RPE-1 cells (C= control, K= oncogenic KRAS OE). HSP90 is used as a loading control. D. Doubling times of parental cells and early clones with and without the expression of exogenous mutant KRAS as determined by live cell imaging. Doubling times were determined 2 weeks after the initial infection with KRAS infected cells once selection was completed. A paired one-way t-test was performed and P-values are assigned according to GraphPad standard. F. Aneuploidy scores were extracted from DepMap and cell lines were categorized on the presence or absence of a KRAS hotspot mutation. An unpaired t-test was performed and P-values are assigned according to GraphPad standard. G. The presence of a chromosome 12p amplification in KRAS WT or mutant cell lines was determined on DepMap data. A Chi-square test was performed to determine the significance of the calculated Odds Ratio. P-values are assigned according to GraphPad standard.

To test whether KRAS overexpression is sufficient to drive adaptation, we infected parental and several early, non-adapted clones with a lentiviral plasmid encoding oncogenic KRAS (DD-HA-PAmCherry-HA-KRAS-G12D (69). We confirmed successful transduction by western blot (Fig. 6C). Strikingly, overexpressing oncogenic KRAS consistently improved proliferation rates in all tested clones, but not in parental cells (Fig. 6D). Importantly, although proliferation rates significantly improved, they did not reach those of fully adapted clones. This suggest that oncogenic KRAS amplification can contribute, but cannot fully drive adaptation to aneuploidy. If oncogenic KRAS mutations contribute to aneuploidy tolerance, we predict that KRAS mutant tumor cell lines have higher aneuploidy levels as they are predicted to be more tolerant. Indeed, we find a significant increase in aneuploidy scores of cell lines harboring such mutation (Fig. 6F). Moreover, Mutant KRAS tumor cell lines are more likely to amplify chromosome 12p, suggesting that enhancing the dose of oncogenic KRAS mutations is a beneficial trait (OR=2.043, p=0.0009).

In summary, our study has demonstrated that human cells possess the capacity to adapt to aneuploidy over time. We show that common aspects of adaptation are the reduction of CIN and the suppression of downstream concurrent inflammation and that one of the drivers of adaptation is the overexpression of oncogenic KRAS.

## Discussion

Aneuploidy is a detrimental condition for untransformed cells, but is notably prevalent in cancer cells. This suggest that cancer cells employ mechanisms to surmount the stress responses associated with aneuploidy, enabling rapid proliferation despite aberrant karyotypes. Here, we aimed to characterize the main cellular changes during adaptation to aneuploidy and uncover the mechanisms underlying this adaptation in human cells.

In accordance with prior findings (2–8), the aneuploid clones we generated initially exhibited reduced proliferation rates. Strikingly, over time all of the clones displayed accelerated growth, often reaching or approaching parental levels, indicative of adaptation to aneuploidy. We found minor karyotype alterations in most clones, mostly towards karyotype simplification. This suggests that a selection for less complex karyotypes can contribute to the process of adaptation. However, it needs to be noted that the simplification was rather modest and a subset of clones even displayed evolved karyotypes resulting in more gene imbalances, while they still displayed improved growth. Intriguingly, in adapted clones, the impact of the number of imbalanced genes on proliferation rates was notably reduced. This implies the involvement of active adaptation mechanisms facilitating better tolerance to gene imbalances.

We observed that adaptation is largely characterized by a correction of the early transcriptomic responses to aneuploidy, suggesting that the early stress responses are limiting growth in response to aneuploidy. Most strikingly, we observed an upregulated inflammatory response in all early clones which was consistently reduced upon adaptation. We show that the reduction of inflammation goes parallel with a reduction in CIN. Intriguingly, a recent paper identified mutations that resulted in the suppression of chromosome missegregations as a key step in adaptation to CIN in yeast (70). This finding supports the notion that CIN exerts a toxic effect on cells and its correction is a favorable way to facilitate fast proliferation. We speculate that the inflammation as a consequence of CIN contributes to this toxicity. To support this, we showed that the transcriptomic changes in early aneuploid clones, including the inflammatory responses, reflect those of parental cells experiencing acute CIN induced by mitotic checkpoint inhibition ((40) and Fig S4A). We show that inflammation as a consequence of acute CIN is complex which can be explained by the fact that CIN-induced inflammation can be prompted by multiple effects. For example, directly via micronuclear rupture or the formation of chromatin bridges resulting in cGAS/STING signaling (41–44), but also more indirectly through NF-kB and expression of SASP genes upon cellular arrest (29,45). STING-depletion was not sufficient to overcome the growth defect of early aneuploid clones. It is possible that we did not suppress cGAS/STING signaling sufficiently or that the duration of the knockdown was insufficient to see an effect. However, it is likely that other aspects of inflammation might be more dominant in driving slow growth. It will be challenging to fully alleviate the observed inflammatory responses, due to the complex nature of CIN-induced inflammation. However, this would be a powerful approach to uncover if and which aspects of the inflammatory response contribute to the slow growth phenotype observed in the early aneuploid clones. Additionally, while correcting CIN in early clones could provide a method to reduce inflammation, this is technically challenging, and previous attempts to do so have been unsuccessful (40). Nevertheless, we provide evidence that both CIN and inflammation are consistently reduced upon adaptation, and that these features are strongly linked to each other.

To understand the mechanism through which cells reduce CIN rates upon adaptation, it is important to gain more insights into the initial responses to aneuploidy that might be responsible for inducing CIN. Previously, we linked the initiation of CIN to proteotoxic stress arising as a consequence of chromosomal gains in early aneuploid clones (40). In accordance with this, in our current study we observed that critical players of the UPR PERK-pathway, ATF3 and CHOP, were significantly upregulated in early clones compared to parental cells, which were corrected to near-parental levels upon adaptation. Activation of PERK is aimed at reducing translational loading of the ER by reducing the global levels of translation (61,71). Consistent with this, we observed a reduction of processes related to translation, peptide biosynthesis, and RNA processing in the early aneuploid clones, responses that have been reported in other aneuploid models as well (72–74). How these alterations related to proteotoxic stress translate into CIN awaits further investigation.

Earlier studies have shown that alleviating proteotoxic stress can facilitate increased proliferation, via deletion of the deubiquitinase Ubp6 in yeast (12,13,20), or via overexpression of HSF1 in human aneuploid cells relieving HSP90-mediated protein folding deficiency (19). These findings underline that the resolution of proteotoxic stress plays an important role in the adaptation process. Besides, other aneuploidy-tolerating mutations have been linked to metabolic rewiring (58,60). Also, aneuploid cells have been shown to exhibit heightened glucose uptake (4). Therefore, we speculate that the reduction in proteotoxic stress observed upon adaptation may be attributed to enhanced energy production, possibly established via metabolic rewiring. Transcriptome analysis indeed suggests an upregulation of glycolysis during adaptation, implying that enhanced glycolysis might be critical in overcoming the negative effects of aneuploidy.

When evaluating large karyotype alterations, we observed a specific karyotype evolution pattern in a subset of our clones upon adaptation. This pattern concerned the gain of 12p or the entire chromosome 12 encoding oncogenic KRAS. Many studies show that specific aneuploidies can be beneficial under conditions of stress (52,53,64,66,75) and the observed 12p amplification is example of a recurrent pattern that emerges in response to aneuploidy-induced stress. Our findings show that increased expression of mutant KRAS is sufficient to drive adaptation to aneuploidy in early clones. A similar gain of chromosome 12 containing mutant KRAS was also described to occur in xenograft-derived aneuploid HCT116 cells (66), indicating that this is an event not exclusively observed in RPE-1 cells. Given the established role of KRAS in metabolic rewiring (76), we speculate that increasing the dose of oncogenic KRAS in our cells promotes adaptation to aneuploidy by enhancing glycolytic rates and thereby overcoming proteotoxic stress. Interestingly, a recent study showed enhanced sensitivity of aneuploid RPE-1 clones to inhibitors of the RAF/MEK/ERK pathway (77). It would be relevant to test if metabolic alterations play a role in this enhanced dependency.

Collectively, our data supports the following model (depicted in Fig. 7): as a consequence of proteotoxic stress, early aneuploid clones are highly CIN resulting in an inflammatory response and reduced proliferation. Over time, cells adapt to aneuploidy, consistently displaying reduced CIN levels and reduced inflammation, which ultimately promotes increased proliferation rates. This is a result of decreased stress responses, potentially driven by metabolic rewiring for example facilitated by oncogenic KRAS amplification. To gain further insights, comprehensive analyses such as full genome sequencing and proteome analysis of the adapted clones could unveil additional pathways required for adaptation. It is tempting to speculate that such alterations aim at relieving proteotoxic stress, possibly by altering the metabolic state of the cell.

**Figure 7:**
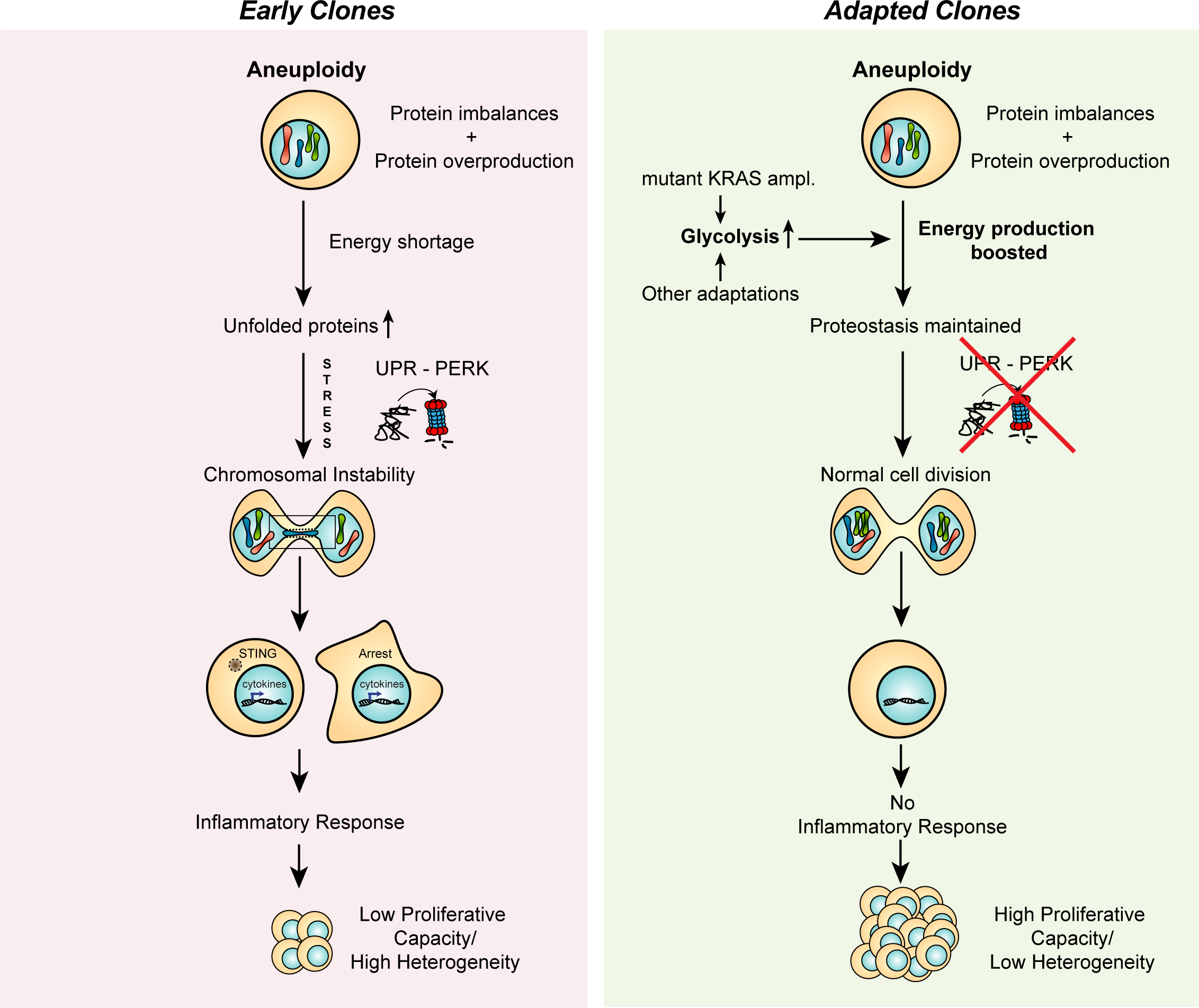
Model A. Early aneuploid cells have problems coping with the extra demands on the protein turnover machineries for the production, folding and degradation of the extra proteins encoded on the aneuploid chromosomes. This results in an excess of unfolded proteins, proteotoxic stress response and potentially additional stress responses. The aneuploidy associated stresses lead to enhanced chromosome segregation errors. These segregation errors lead to inflammatory signaling via the cGAS-STING pathway and/or via SASP-like signaling. These phenotypes contribute to the slow proliferation associated to aneuploidy. Upon adaptation, aneuploidy induced stresses are reduced, likely due to metabolic rewiring such as enhanced glycolysis. Oncogenic KRAS amplifications could contribute to metabolic rewiring but other mechanisms can drive such a switch as well. The restored energy production and reduced stress signaling prevent chromosome missegregations and inflammation and are therefore compatible with high proliferative capacity.

## Acknowledgements

We thank the Genomics Core Facility of the Netherlands Cancer Institute for sample preparation, data acquisition, and analysis of CNV and RNA sequencing experiments. This work was funded by a KWF Young Investigator Grant – 12233 to J. Raaijmakers.

## Author contributions

R.H.M. and J.A.R. conceived the project. D.C.H., and J.A.R. performed and analyzed the experiments. M.S. performed data processing and data analysis. D.C.H., J.A.R., and R.H.M. wrote the manuscript.

## Materials and Methods

### Cell culture, cell lines, and reagents

hTert-immortalized retinal pigment epithelium (RPE-1) p53kd cells were kindly provided by R. Beijersbergen. RPE-1 p53kd cells were generated by transduction with pRetroSuper-p53 (with the shRNA sequence 5’- CTACATGTGTAACAGTTCC-3’) and selected with Nutlin-3a for functional loss of p53. H2B-Dendra2 cells were made as described in (55). Cells were cultured at 37C at 5% CO2 in Advanced Dulbecco’s Modified Eagle Medium: Nutrient mixture F-12 (DMEM-F12) with Glutamax (GIBCO), supplemented with 12% FCS (Clontech), 100 U/ml penicillin (Invitrogen), 100 μg/ml streptomycin (Invitrogen) and 2mM UltraGlutamin (Lonza). Inhibitors were all dissolved in DMSO and were used at the following concentrations: Mps1 inhibitor (NMS- P715) (480uM) and a CENP-E inhibitor (GSK923295) (50nM).

### Generating aneuploid clones

Clones were generated by blocking RPE-1 p53kd cells in Thymidine for 14 hours, after which they were released in medium containing a combination of an Mps1 inhibitor (NMS-P715) (480uM) and a CENP-E inhibitor (GSK923295) (50nM) for 8 hours to induce whole chromosome aneuploidies and segmental aneuploidies a shown in(55). After treatment, cells were collected by trypsinization and cells were plated single cell in 384-well plates. On the same day, wells were examined for the presence of individual cells to ensure a single cell was present. With this approach we successfully generated 14 aneuploid clones (Fig. S1A). Some of these clones were described in a previous study (40). The gain of 10q in parental RPE-1 cells, deriving from an imbalanced fusion of the q-arm of chromosomes 10 to the X chromosome ((78), ATCC) was not considered as a *de novo* aneuploidy (Fig. S1).

### Live-cell imaging

For live-cell imaging, cells were grown in a Lab-Tek II chambered coverglass (Thermo Science). Images were acquired every 5 minutes using a DeltaVision Elite (Applied Precision) microscope maintained at 37°C, 40% humidity and 5% CO2, using a 20x 0.75 NA lens (Olympus) and a Coolsnap HQ2 camera (Photometrics) with 2 times binning. DNA was visualized using SiR-DNA (Spirochrome, 0.25μM) or using SPY-650 (Spirochrome, diluted according to manufacturer, used 1:5000). Image analysis was done using ImageJ software and all conditions were blinded before analysis.

### Cell growth analysis

Proliferation was measured by using a Lionheart FX automated microscope (Biotek). For these experiments 500 cells were plated in 96-well plates. Two or three replicate wells were imaged per clone with a 4 h interval for 5 days, and cells were stained with the DNA dye siR-DNA (Spirochrome, 0.25μM) or using SPY-650 (Spirochrome, diluted according to manufacturer, used 1:5000). Proliferation rates were measured by performing cell count analysis using Gen5 software (BioTek) and doubling times were calculated using GraphPad Prism 8 software during exponential growth phase.

### Immunoblotting

RPE-1 cells were harvested and lysed using Laemmli buffer (120 mM Tris, pH 6.8, 4% SDS, and 20% glycerol). Equal amounts of protein were separated on a polyacrylamide gel and subsequently transferred to nitrocellulose membranes. Membranes were probed with the following primary antibodies (1:1000): pan-RAS (Rabbit, Cell Signaling, #3965), HSP90 α/β (rabbit, Santa Cruz, sc-13119). HRP-coupled secondary antibodies (Dako) were used in a 1:1000 dilution. The immunopositive bands were visualized using ECL Western blotting reagent (GE Healthcare) and a ChemiDoc MP System (Biorad).

### Copy number analysis

DNA was isolated using the DNeasy Blood and Tissue kit (Qiagen) according to the manufacturer’s protocol. The number of double-stranded DNA in the genomic DNA samples was quantified by using the Qubit dsDNA HS Assay Kit (Invitrogen, cat no Q32851). Up to 2000 ng of double-stranded genomic DNA was fragmented by Covaris shearing to obtain fragment sizes of 160-180 bp. Samples were purified using 1.8X Agencourt AMPure XP PCR Purification beads according to the manufacturer’s instructions (Beckman Coulter, cat no A63881). The sheared DNA samples were quantified and qualified on a BioAnalyzer system using the DNA7500 assay kit (Agilent Technologies cat no. 5067-1506). With an input of maximum, 1 μg sheared DNA, library preparation for Illumina sequencing was performed using the KAPA HTP Library Preparation Kit (KAPA Biosystems, KK8234). During library enrichment, 4-6 PCR cycles were used to obtain enough yield for sequencing. After library preparation, the libraries were cleaned up using 1X AMPure XP beads. All DNA libraries were analyzed on a BioAnalyzer system using the DNA7500 chips to determine the molarity. Up to eleven uniquely indexed samples were mixed by equimolar pooling, in a final concentration of 10nM, and subjected to sequencing on an Illumina HiSeq2500 machine in one lane of a single read 65 bp run, according to manufacturer’s instructions.

Low-coverage whole-genome samples, sequenced single-end 65 base pairs on the HiSeq 2500 were aligned to GRCh38 with bwa version 0.7, mem algorithm (79). The mappability per 15 kilobases on the genome, for a sample’s reads, phred quality 37 and higher, was rated against a similarly obtained mappability for all known and tiled 65bp subsections of GRCh38; a reference genome based mappability provided by QDNAseq (80), using a GRCh38 lifted version (https://github.com/asntech/QDNAseq.hg38.git). QDNAseq segments data using an algorithm by DNAcopy (81) and calls copy number aberrations using CGHcall (82), and visualization was adapted from the QDNAseq code.

### TIDE

TIDE (tracking of indels by decomposition) was used to estimate the ratio of WT KRAS to mutant KRAS shown in Fig. 6A. In short, the region harboring the 6 base pair insertion was amplified via PCR following primers: forward primer: CTTAAGCGTCGATGGAGGAG and reverse primer: TGTATCAAAGAATGGTCCTGCAC. The PCR products were treated with exonuclease I (Biolabs, M0293S) and were subjected to Sanger Sequencing using the forward primer and analyzed by the TIDE method (68), using WT HeLA cells as a reference, as these cells harbor WT KRAS.

### Determining the number of imbalanced genes

For clones harboring large chromosome aneuploidies, the number of imbalanced/gained/lost coding genes was calculated by first determining gained and lost segments per clone using RNA sequencing data. A chromosomal segment was considered a gain if the average expression of genes on that segment passes a threshold of 0.31 log2 fold change (corresponding to a 1.25-fold overexpression). A segments was considered a loss if the average expression of genes on that segment was below a threshold of -0.415 log2 fold change (corresponding to 0.75-fold expression). Then, per clone the number of coding genes located on the gained or lost segments were summed up to obtain the number of lost coding or gained genes respectively. The lost and gained coding genes were added up to obtain the total number of imbalanced gene per clone.

### RNA sequencing and data analysis

RPE-1 p53KD cells (parental and clones) were harvested in buffer RLT (Qiagen). Strand-specific libraries were generated using the TruSeq PolyA Stranded mRNA sample preparation kit (Illumina). In brief, polyadenylated RNA was purified using oligo-dT beads. Following purification, the RNA was fragmented, random-primed and reverse transcribed using SuperScriptII Reverse Transcriptase (Invitrogen). The generated cDNA was 3′ end- adenylated and ligated to Illumina Paired-end sequencing adapters and amplified by PCR using HiSeq SR Cluster Kit v4 cBot (Illumina). Libraries were analyzed on a 2100 Bioanalyzer (Agilent) and subsequently sequenced on a Nova Seq 6000 (Illumina). We performed RNAseq alignment using TopHat 2.1.1. on GRCh38 and counted reads using Rsubread 2.4.3 (Ensembl 102).

We calculated differential expression between two biological replicates of the parental and each clone, as well as the Mps1 and CENP-E inhibitor treated vs control, using DESeq2 1.31.3. Copy numbers from RNA-seq data were determined using a Generalized Additive Model (GAM) smoothing (mgcv R package) with a gamma parameter of 2 and a weight parameter of 1/lfcSE. We tested for gene set differences by using a linear regression model of the Wald statistic (as reported by DESeq2) between genes belonging to a set vs. genes not belonging to a set. Gene set collections included MSigDB hallmarks (2020) and Gene Ontology (2021). For the gene sets related to inflammation, we used manually generated gene lists based on published literature. For STING signaling we used gene lists generated by Santaguida et al. 2017 (29), as well as a list of genes that was shown to be at least 2,5-fold differentially expressed with a P-value<0.05 upon treatment with cGAMP in primary human cells (83). For NFkB signaling we took a list of known NFkB-target genes published by the Boston University (https://www.bu.edu/nf-kb/gene-resources/target-genes/). For the SASP signaling, we used a recently published signature (84). See Table S1 for the different gene sets.

### RNA isolation and qRT-PCR analysis

Total RNA was extracted from untreated RPE-1 cells. RNA isolation was performed by using the Qiagen RNeasy kit and quantified using NanoDrop (Thermo Fisher Scientific). cDNA was synthesized using SuperScript III reverse transcription, oligo dT (Promega), and 1000 ng of total RNA according to the manufacturer’s protocol. Primers were designed with a melting temperature close to 60 degrees to generate 90–120-bp amplicons, mostly spanning introns. cDNA was amplified for 40 cycles on a cycler (model CFX96; Bio-Rad Laboratories) using SYBR Green PCR Master Mix (Applied Biosystems). Target cDNA levels were analyzed by the comparative cycle (Ct) method and values were normalized against β-actin expression levels.

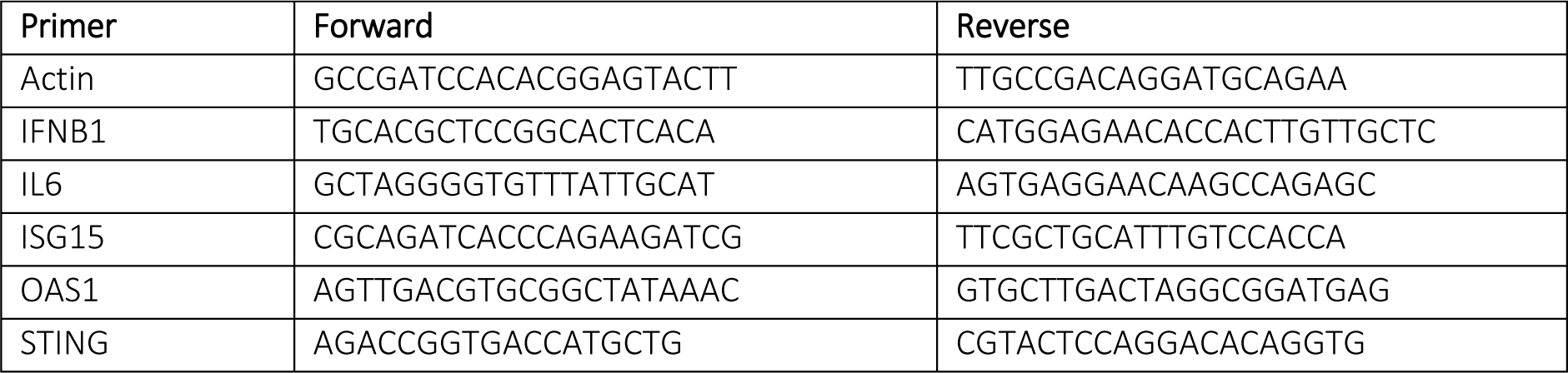

### siRNA and crRNA transfections

siRNA transfections were performed using RNAiMAX (Invitrogen) according to the manufacturer’s guidelines. The following siRNAs were used in this study: siNT (Non-targeting; Dharmacon), siTMEM173 (Dharmacon, OTP- SMARTpool). SiRNAs were used at a final concentration of 20μM.

## Supplementary Figure Legends

**Supplementary Figure 1:**
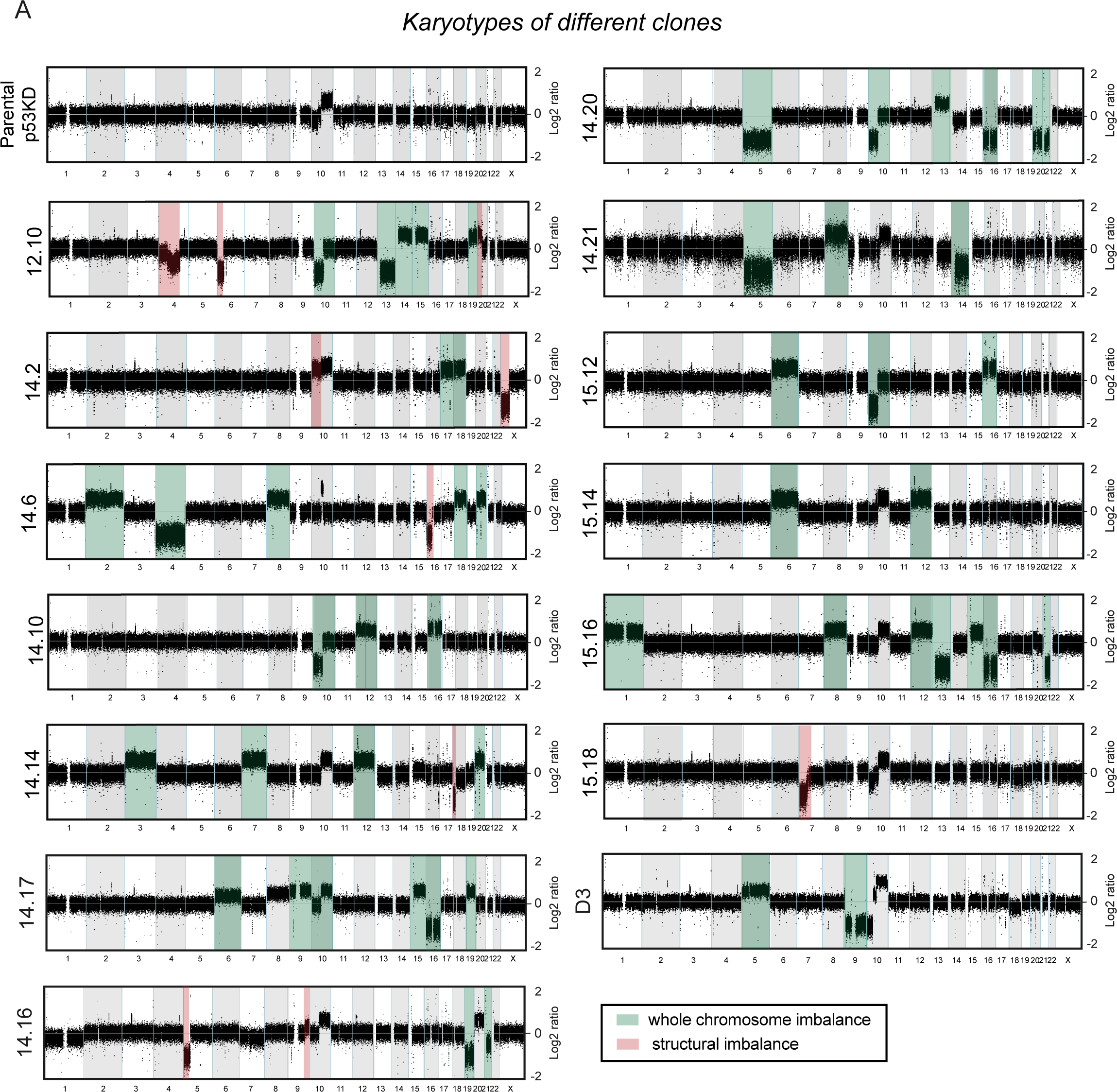
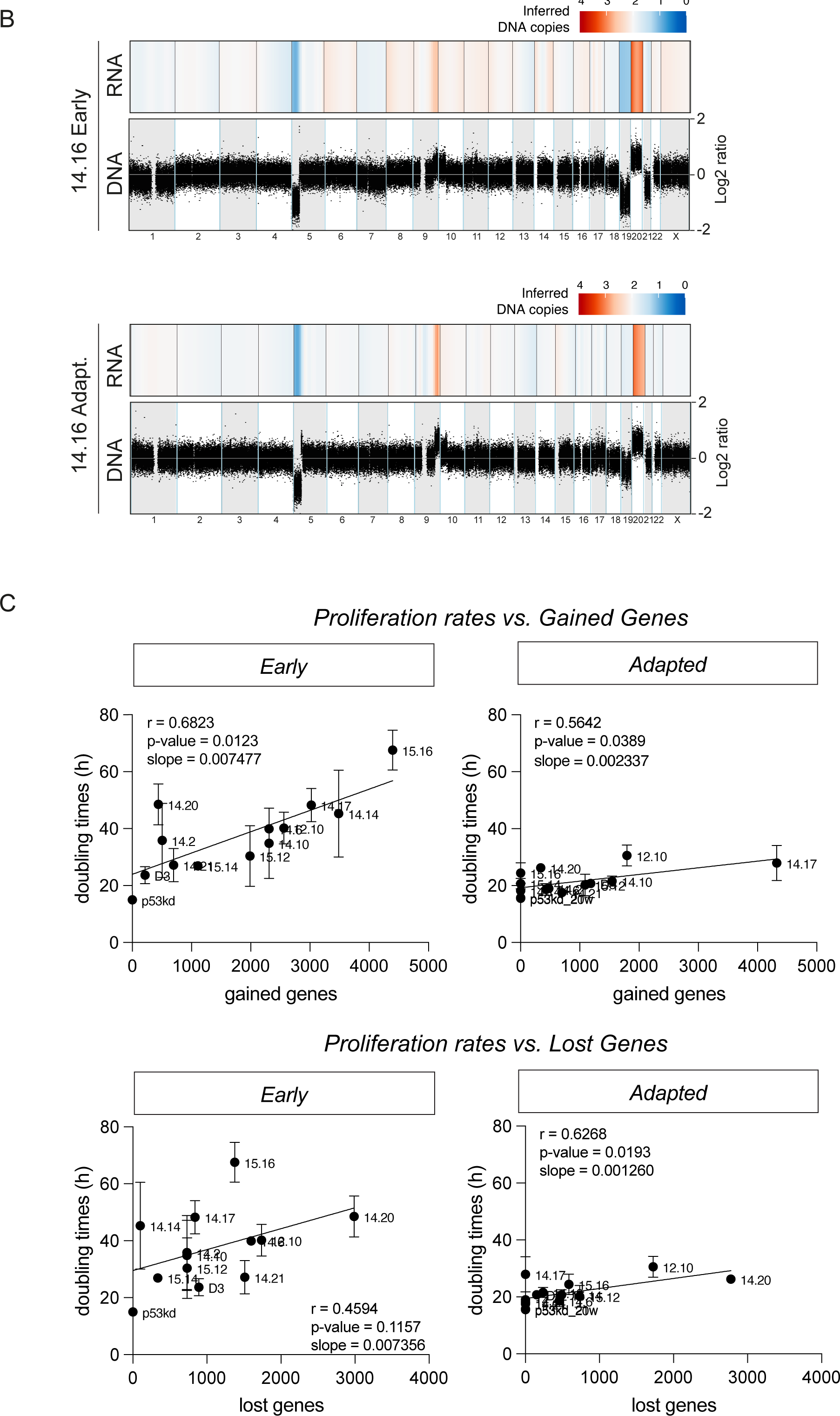
Copy number analysis and correlation between proliferation and gene imbalances A. Genome-wide chromosome copy number profile as determined by CNV-seq of the RPE-1 p53kd parental clone and 14 clones harboring chromosome imbalances. Chromosome gains and losses were depicted in green boxes (whole chromosomes) or red boxes (chromosome segments). Alterations of chromosome 10, already present in the parental cells, were not highlighted. B. Comparison between RNA and DNA sequencing for inferring karyotypes for clone 14.16, both early passage and adapted. Both RNA read counts and DNA read count were normalized over parental early. DNA copy number is displayed in Log2 ratio per bin and RNA sequencing are displayed as inferred copy number per bin. C. Spearman correlation between RNA sequencing-derived gained and lost genes per clone and proliferation rates measured in Fig. 1B for early and adapted clones Error bars indicate standard deviation. Line shows linear regression model fitted to the data.

**Supplementary Figure 2:**
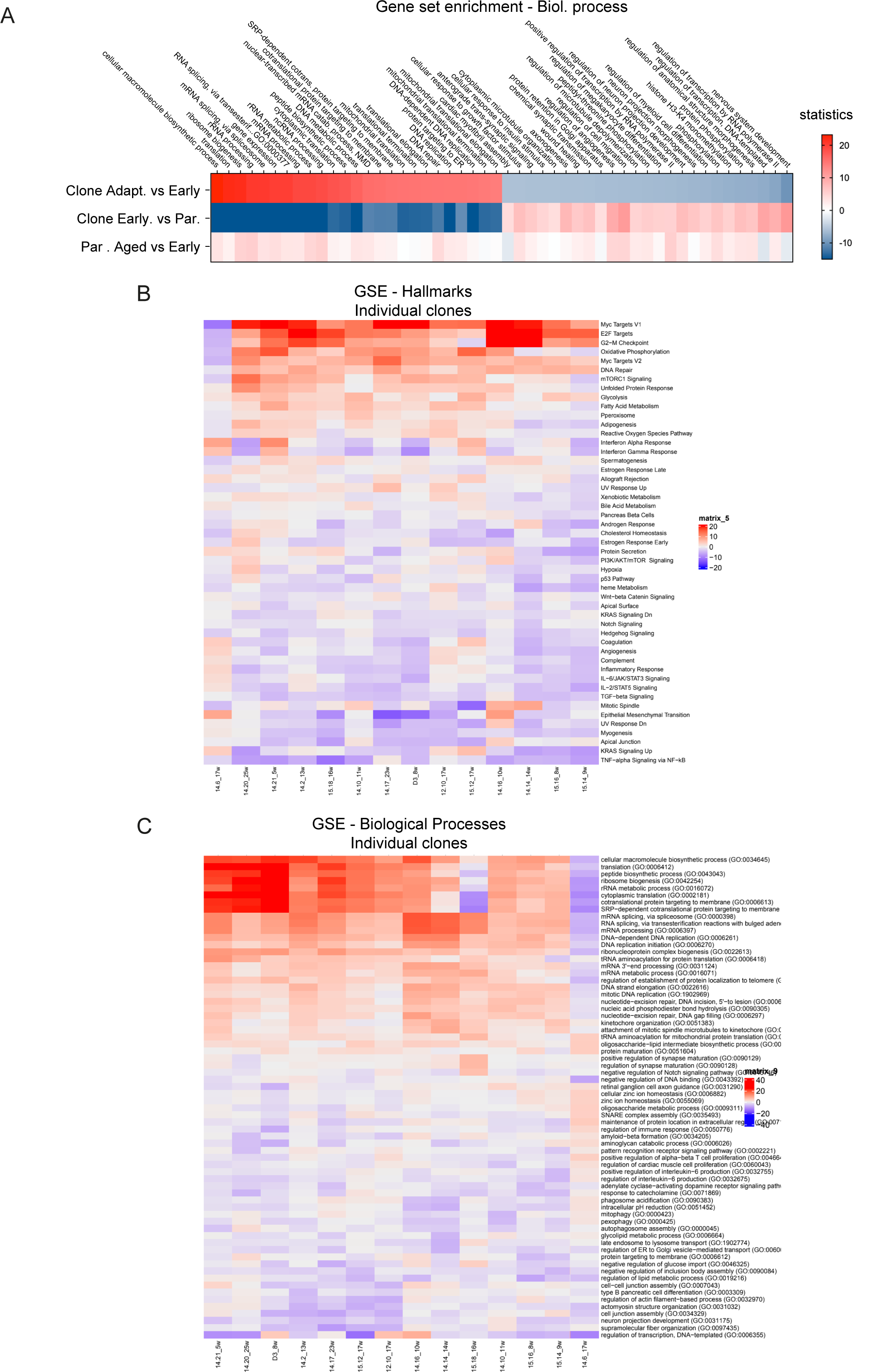
Gene set enrichment for biological processes and individual clones A. Gene set enrichment analysis for GO Biological processes on the transcriptional alterations observed in all adapted clones compared to their early counterparts. The top 50 most altered gene sets are displayed (top row). Alterations in early clones over parental (middle row) and aged parentals (bottom row) are displayed as a reference. Colors indicate Wald statistics. B. Gene set enrichment analysis (GSEA) for the individual clones, evaluating up and downregulated GO hallmarks in adapted clones compared to their early counterparts. Two replicates of every clone were sequenced. Colors indicate Wald statistics. C. Gene set enrichment analysis (GSEA) for the individual clones, evaluating up and downregulated GO biological processes in adapted clones compared to their early counterparts. Two replicates of every clone were sequenced. Colors indicate Wald statistics.

**Supplementary Figure 3:**
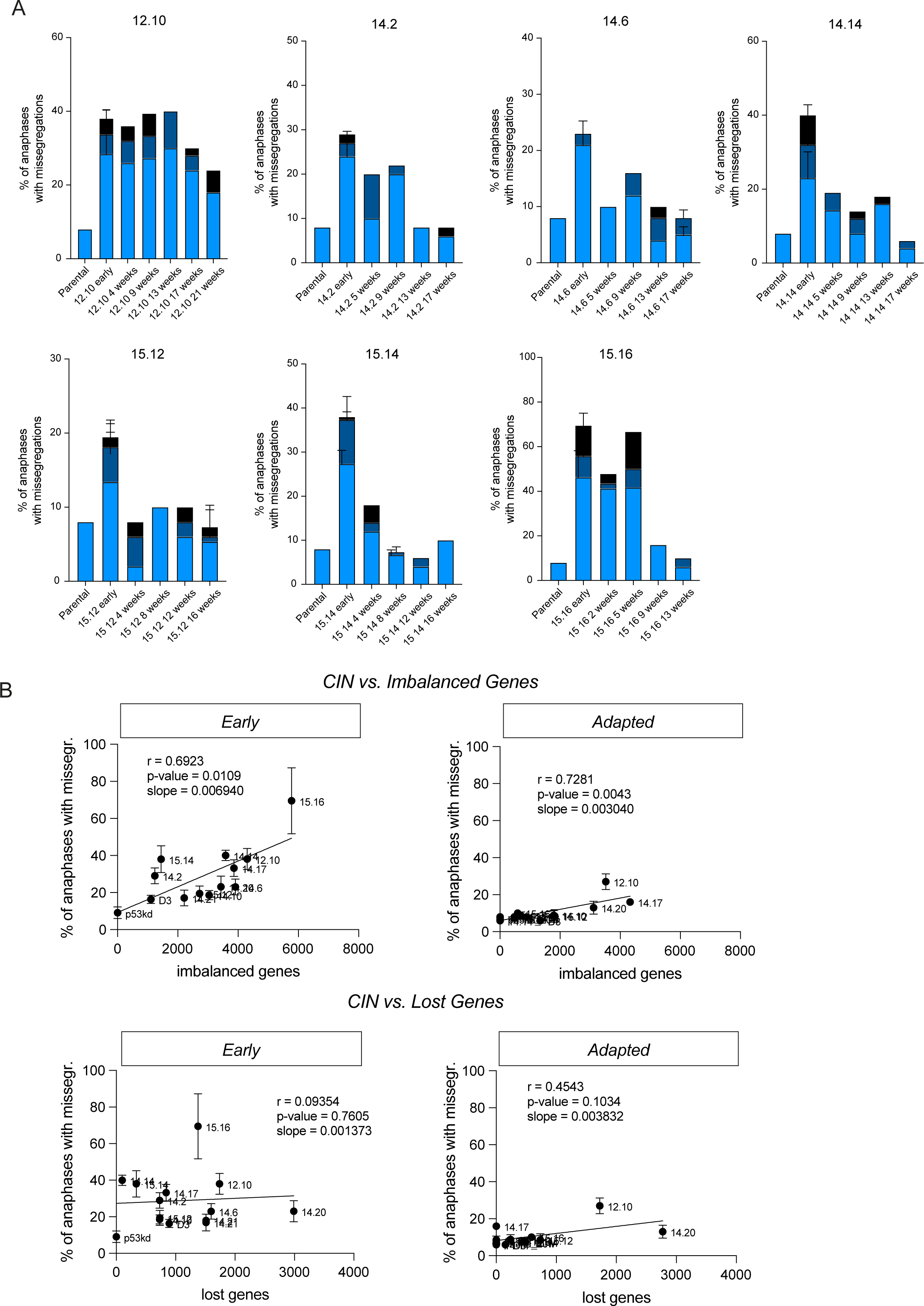
CIN is corrected during the trajectory of adaptation A. Chromosome missegregation rates determined at monthly interval by live cell imaging of parental RPE-1 p53KD cells and aneuploid clones at from Fig. S1A, divided into three subcategories: lagging chromosomes, anaphase bridges and others (multipolar spindle, polar chromosome, cytokinesis failure, binucleated cell). All conditions were analyzed blinded. Bars are averages of at least 2 experiments and a minimum of 50 cells were filmed per clone per experiment. Error bars indicate standard deviation. An ordinary one-way ANOVA was performed between parental and clones. P-values are assigned according to GraphPad standard. B. Spearman correlation between the number of RNA-sequencing derived imbalanced and lost genes per clone and the level of CIN as percentage of total number of anaphases as determined in Fig. 4A. Error bars indicate standard deviation.

**Supplementary Figure 4:**
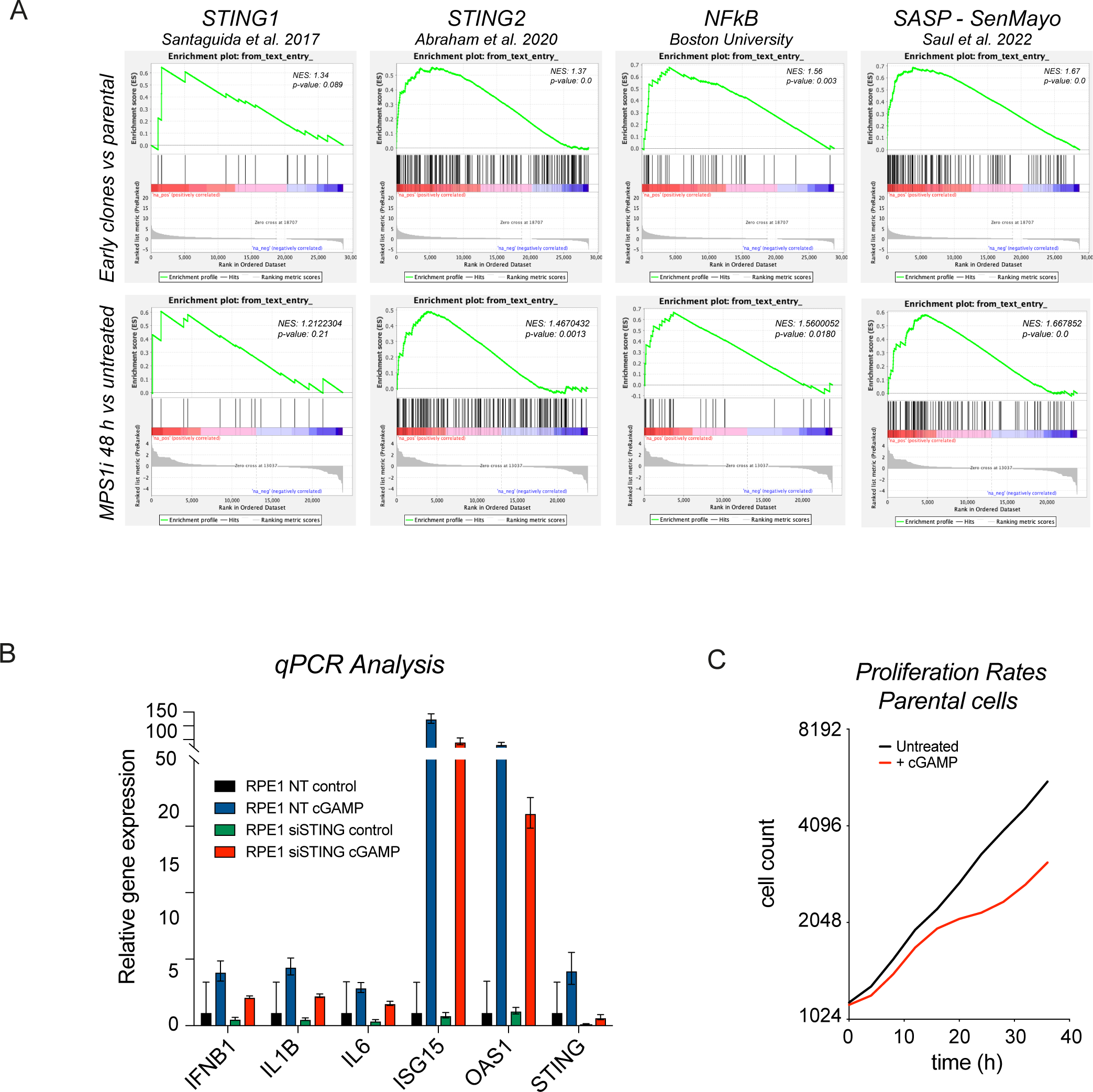
Characterization of the inflammatory response A. Transcriptome data of early clones (top row) and of parental p53KD cells treated for 48h with 50nM CENP- Ei and 480nM MPS1i tested against known STING signaling, NF-κB signaling and SASP signaling gene sets. An overview of the different gene sets can be found in table S1. B. mRNA levels of inflammatory response cytokines determined via qRT-PCR of parental RPE-1 p53KD after 72 hours siRNA against STING, untreated or treated for 24 hours with 67μM of cGAMP. Values were normalized to actin and are displayed relative to expression levels in untreated parental cells. Bars show mean expression levels; error bars indicate upper and lower limits. B. Cell counts of parental RPE-1 p53KD cells left untreated or treated with 67μM cGAMP which was added 2 hours prior to imaging. Y-axis Is displayed in a Log2 scale.

